# The gill-associated symbiont microbiome is a main source of woody-plant polysaccharide hydrolase genes and secondary metabolite gene clusters in *Neoteredo reynei*, a unique shipworm from south Atlantic mangroves

**DOI:** 10.1101/357616

**Authors:** Thais L. Brito, Amanda B. Campos, F.A. Bastiaan von Meijenfeldt, Julio P. Daniel, Gabriella B. Ribeiro, Genivaldo G. Z. Silva, Diego V. Wilke, Daniela Morais, Bas E. Dutilh, Pedro M. Meirelles, Amaro E. Tridade-Silva

**Affiliations:** Drug Research and Development Center, Department of Physiology and Pharmacology, Federal University of Ceara, Fortaleza, CE, Brazil; Institute of Biology, Federal University of Bahia, Salvador, BA, Brazil; Theoretical Biology and Bioinformatics, Utrecht University, Utrecht, Netherlands; Computational Science Research Center, San Diego State University, San Diego, CA, USA; National Institute of Metrology, Quality and Technology, Rio de Janeiro, Brazil; Centre for Molecular and Biomolecular Informatics, Radboud University Medical Centre, Nijmegen, The Netherlands.

**Author notes:** Corresponding author (AETS).

## Abstract

Teredinidae is a family of highly adapted wood-feeding and wood-boring bivalves, commonly known as shipworms, whose evolution is linked to the acquisition of cellulolytic gammaproteobacterial symbionts harbored in bacteriocytes within the gills. In the present work we applied metagenomics to characterize microbiomes of the gills and digestive tract of *Neoteredo reynei*, a mangrove-adapted shipworm species found over a large range of the Brazilian coast. Comparative metagenomics grouped the symbiotic gammaproteobacterial community of gills of different *N. reynei* specimens, indicating closely related bacterial types are shared, while intestine and digestive glands presented related, and more diverse microbiomes that did not overlap with gills. Annotation of assembled metagenomic contigs revealed that the symbiotic community of *N. reynei* gills was a hotspot of woody-polysaccharides degrading hydrolase genes, and Biosynthetic Gene Clusters (BGCs), while in contrast, the digestive tract microbiomes seems to play little role in wood digestion and secondary metabolites biosynthesis. Metagenome binning recovered the nearly complete genome sequences of two symbiotic *Teredinibacter* strains from the gills, a representative of *Teredinibacter turnerae* “clade I” strain, and a yet to be cultivated *Teredinibacter* sp. type. These *Teredinibacter* genomes, as well as unbinned gill-derived gammaproteobacteria contigs, code for novelty including an endo-β-1,4-xylanase/acetylxylan esterase multi-catalytic carbohydrate-active enzyme, and a trans-acyltransferase polyketide synthase (trans-AT PKS) gene cluster with the gene cassette for generating β-branching on complex polyketides. Multivariate analyzes have shown that the secondary metabolome encoded on the genomes of *Teredinibacter* representatives, including the genomes binned from *N. reynei* gill’s metagenomes, stand out within the Cellvibrionaceae family by size, and enrichments for polyketide, nonribosomal peptide and hybrid BGCs. Results grouped here add to the growing characterization of shipworm symbiotic microbiomes and indicate that the *N. reynei* gill gammaproteobacterial community is a prolific source of biotechnologically relevant enzymes for wood-digestion and bioactive compounds production.

## Introduction

Teredinidae, a family of obligate xylotrepetic (wood-boring) and xylotrophic (wood feeding) mollusks, evolved from an ancestor lineage within the eulamellibranch order Myoida, and succeeded in exploiting wood as new food source, most probably by concomitantly acquiring cellulolytic bacterial symbionts [1]. These animals suffered dramatic body shape adaptations to thrive as wood feeders including greatly reduced and burrowing-specialized valves (shells), and a wood-sheltered worm-like body plan protruding from their shells, which led to their common name of shipworms [2].

Shipworm bacterial symbiotic communities are formed by cultivated and as yet uncultivated closely related gammaproteobacteria, located intracellularly in bacteriocytes within the gills (ctenidia) tissue [3–6]. The prototypical shipworm symbiont is the cellulolytic and nitrogen fixing bacterium *T. turnerae*, isolated from a great variety of Teredinidae species from all over the globe [7,8]. Sequencing of the complete genome of one strain of *T. turnerae*, T7901, revealed that the genome of this bacterium codes for an arsenal of enzymes specialized in breaking down woody material, reinforcing the hypothesis that shipworm gammaproteobacterial symbionts play a critical role on supporting host nutrition[9]. Indeed, two seminal works provided strong evidence for this hypothesis: firstly multi isotopic mass spectroscopy (MIMS) combined with transmission electron microscopy (TEM) showed, in *Lyrodus pedicellatus*, that *T. turnerae* cells could fix atmospheric N_2_ *in symbio*, and then transfer nitrogenized compounds to their host [10]. Secondly, a combination of metagenomics and proteomics of *Bankia setacea*, showed that wood-specialized hydrolytic enzymes secreted by the symbiotic community are, somehow, selectively conducted from the gill symbiotic site up to the host‘s cecum to reach their substrates, the ingested wood particles [11].

Characterization of the *T. turnerae* T7901 genome also revealed that this endosymbiotic bacterium contains a diversity of Biosynthetic Gene Clusters (BGCs) for secondary metabolites comparable to those observed in free-living *Streptomyces* species known for their proficiency at secondary metabolite production[9]. Indeed, compounds of the tartrolon family of boronated antibiotic [12], and a novel triscatecholate siderophore called turnerbactin [13], were purified from chemical extracts of *T. turnerae* cultures and had their respective biosynthetic routes identified and characterized by retrobiosynthetic and phylogenetic approaches. Moreover, both BGCs were proven to be expressed *in symbio* and signatures of these compounds were detected by mass spectroscopy of shipworms whole animal extracts [12,13]. Such results fueled a hypothesis that bioactive secondary metabolites could also play a role in supporting the symbiosis by providing chemical defense and so, improving holobiont fitness [14], and/or mediate competition/specialization to shape the distribution of bacterial types within the gills. Fluorescence *in situ* hybridization (FISH) and confocal microscopy surveys of tissues from several shipworm species provided evidence supporting both these hypothesis, showing that i) symbiotic gammaproteobacterial types are structured within the gills, in separated niches of bacteriocytes, and ii) tissues of the shipworm digestive system are virtually sterile (cecum) or containing a discrete microbial community (intestine) [11,15].

Recently, the gill symbiotic community of the giant shipworm *Kuphus polythalamia,* a rare species that burrows into mud and sediment instead of wood, was shown to be composed of sulfur oxidizing chemoautotrophic gammaproteobacteria instead of Teredinibacter-related cellulolytic types [16]. These findings suggested this bivalve is a chemoautotrophic descendent of a xylotrophic ancestor, thus adding complexity to the biology of symbiosis in the family. *N. reynei* is another singular shipworm that has been reported on mangroves of the south Atlantic, including records of a large range of the 7.367 km long Brazilian coastline, from the States of Pará, at the north of the country, to Santa Catarina, at the very south. *N. reynei* is the only species of the genus, and can be rapidly identified by a unique feature: the presence of large and highly vascularized dorsal lappets at the posterior end of the animal‘s body, whose function is yet to be clarified [2,17]. *N. reynei* is a basal member of the family and has, characteristically, unsegmented pallets, a globular (type II) stomach and an intestine that makes a loose loop forward embracing the crystalline style-sac before running backwards toward the cecum [1]. The *N. reynei* digestive tract includes also two digestive glands, one previously attributed to wood-digestion, another attributed to filter feeding digestion, and both connected to the style-sac through ducts. In addition, *N. reynei* possesses a large cecum, which can reach up to 44% of the animal‘s body size, and an enormous and cylindrically shaped anal canal whose aperture is controlled by a strong muscular sphincter [18]. *N. reynei* gills extend from the base of the siphons up to the posterior end of the visceral mass as demibranchs and then reduce to a food groove at the anterior end of the animal‘s body, without presenting an anterior gill, like in most other shipworm species. Therefore, it comprises only 20% of *N. reynei* body length, a much shorter proportion when compared to other shipworm species. Such reduced gills combined with a large cecum and anal canal, both continuously filled with wood particles, were taken as anatomical evidences that *N. reynei* diet is primarily wood [2,17,18].

Previous culturing efforts lead to isolation of *T. turnerae* strains from *N. reynei* gill homogenate, including CS30, a strain that displayed antimicrobial activity against a spectrum of gram positive and negative indicator bacteria [19]. However, to date, the diversity and biotechnological potential of *N. reynei* gill symbiotic microbiome was still to be explored by culture independent methods.

In the present work we applied metagenomic analysis to explore the *N. reynei* gill symbiotic bacterial community, as well as the microbiome of the digestive glands and intestine, as representative tissues of the shipworm digestive tract. Two symbiotic bacterial genome bins were recovered and analyzed for their capability for producing hydrolytic enzymes and bioactive secondary metabolites. Our findings reinforced the importance of the shipworm symbiotic system as a subject of study and discovery.

## Material and Methods

### Specimens

Three whole adult specimens of *N. reynei* shipworm were collected, using care to avoid injury, from decaying wood chunks in the Coroa Grande mangrove area at Sepetiba Bay, Rio de Janeiro State, Brazil (22.91°S, 43.87°W) on December 5, 2014, and transported to the Laboratory of Biotechnology at the National Institute of Metrology, Quality and Technology (Inmetro, RJ) in sterilized glass jars.

### Metagenomic DNA extraction, sequencing, annotation, and statistics

Digestive glands, intestines and gill tissues were dissected from freshly collected animals (out of wood for ~2hs at 24^°^C) snap frozen and ground in liquid nitrogen, and processed for metagenomic DNA extraction using 2% CTAB lysis buffer, phenol-chloroform-isoamyl alcohol (25:24:1) deproteinization, and the columns from PowerSoil^®^ DNA Isolation Kit (MO BIO Laboratories, Carlsbad, CA,USA) for final purification, as previously described [20]. For each specimen, the two digestive glands were combined prior to grounding for DNA purification. Additionally, two independent DNA purifications were performed from pulverized intestine and gills tissues (replicates), totalizing 5 metagenomic DNA samples per shipworm specimen. Metagenomic DNA samples purity and integrity were respectively evaluated by quantification with a NanoDrop spectrophotometer (Thermo Fisher Scientific Inc.), and by 1% agarose gel electrophoresis. A Qubit fluorimeter (Thermo Fisher Scientific Inc.) was used for accurate double-stranded DNA quantification. Metagenomic DNA libraries were prepared for the fifteen high-quality metagenomic DNA samples, using a *Nextera XT DNA Sample Preparation Kit* (Illumina), as recommended by the manufacturer. Metagenomic libraries were sequenced using the 600-cycle (300 bp paired-end runs) *MiSeq Reagent Kits v3* chemistry (Illumina) with a MiSeq Desktop Sequencer (Illumina) at the Center for Genomics and Bioinformatics (CeGenBio) of the Drug Research and Development Center (NPDM), at the Federal University of Ceara, Brazil. Raw sequence data were submitted to the MG-RAST server (version 4.03) [21] for sequence paired-end joining, quality control, and automated annotation pipeline. Taxonomic annotations were performed using RefSeq, and functional signatures were retrieved with both Clusters of Orthologous Groups (COG) and Subsystems technology databases using the default cutoffs settings (Supplementary Table S1). Taxonomical and functional signatures were submitted to comparative metagenomics using R packages and the Statistical Analysis of Metagenomic Profiles (STAMP) software package, version 2.1.3[22]. Multivariate statistical analyzes were performed with R using the following unsupervised learning techniques: i) hierarchical clustering, with Ward grouping method on a Euclidean distance matrix, ii) PCA biplot, and iii) supervised/unsupervised random forest at R, as previously reported [23]. For hierarchical clustering, cluster significance was accessed by bootstrap resampling using pvclust algorithm[24]. At STAMP, “two groups” comparisons were conducted considering the two main clades obtained by unsupervised hierarchical clustering. For that, the prokaryotic taxonomic (Class level of RefSeq) and functional (Level 2 of the Subsystems technology) features of each group were retrieved from MG-RAST and compared using two-sided Welch‘s t-Test with confidence interval of 0.95 and considering Benjamini-Hochberg False Discovery Rate corrected P-values (q-values) < 10^-5^ as significantly relevant.

### Metagenome assembly, cross-comparison and contig annotation

For annotation-independent cross-assembly comparison between the 15 metagenomic samples generated in this study we used crAss tool [25]. First, all paired-end joined quality controlled metagenome reads of all tissues and specimens were assembled together using SPAdes 3.8 in -meta mode, and k-mer sizes of 21, 33, 55, and 77 [26]. Reads were mapped back to contigs using Bowtie2 [27] and the SAM file was used on crAss, which computes the abundance of reads from each metagenome contributing to form each contig in the cross assembly. The coverage of each of the fifteen metagenomes in contigs was used to build a distance matrix that was displayed in the form of a cladogram [25]. The tissue-specific contigs retrieved from gills, intestine and digestive glands datasets were taxonomically annotated with the contig annotation tool (CAT) [28] which predicts protein-coding genes on a contig, maps these to NCBI’s non-redundant protein database (NR), and subsequently employs a last common ancestor algorithm to give a conservative estimate of taxonomy for the contig. CAT was run with the b1 parameter set to 10, and the b2 parameter set to 0.5. Prodigal 2.6.3 [29] and Diamond 0.9.14 [30] were employed for protein calling and database mapping, respectively. NR was downloaded on November 23, 2017.

### Symbiotic genome binning

Metagenomic sequencing reads were cross-assembled for each tissue with SPAdes 3.8.0 in –meta mode [26]. Tissue-specific contig sets were each binned into draft genome sequences using MetaBAT 0.26.3 [31] that exploits signatures that are typical of contigs belonging to the same genome, including the nucleotide usage and the abundance of the contigs in the different samples. In order to get contig abundances, reads of all the samples were mapped to the tissue-specific assemblies with Burrows-Wheeler Aligner (BWA) 0.7.12 with the BWA-MEM algorithm[32]. Mappings were converted to the binary BAM format with SAMtools 0.1.19 [33] and depth files were created with the script jgi_bam_summarize_contig_depths that is supplied with MetaBAT. Binning with MetaBAT was performed with the sensitive setting, after tests with other presets showed this to give the best results in terms of bin completeness and contamination. Bin completeness, contamination, and strain heterogeneity were assessed using CheckM 1.0.7 [34] based on the presence and absence of sets of single-copy marker genes. CheckM was run in the lineage specific work flow, which places the bins in a reference tree and based on that placement searches for lineage-specific marker genes. Completeness, contamination, and strain heterogeneity were assessed on the lowest level of placement (node) in the CheckM backbone tree. Two high quality genome bins designated gills.bin.1 and gills.bin.4 (>50% completeness and <5% contamination, see Table 1) were generated from the gill assembly, as discussed in the Results section. No high-quality bins could be generated from the digestive gland or intestine assemblies.

**Table 1.** *N. reynei* gill symbiotic genome bins metrics and features.

| Genome bin/Features | Gills.bin.1 | Gills.bin.4 |
| --- | --- | --- |
| **Marker lineage** | s_turnerae (UID4667) | g_Teredinibacter (UID4659) |
| **# genomes** | 2 | 13 |
| **# markers** | 1792 | 1055 |
| **# marker set** | 305 | 271 |
| **Completeness** | 87.12 | 96.45 |
| **Contamination** | 2.8 | 4.88 |
| **Strain heterogeneity** | 0 | 55 |
| **Genome size (bp)** | 4418468 | 5127066 |
| **# contigs** | 355 | 207 |
| **N50 (contigs)** | 16757 | 39993 |
| **Mean contig length (bp)** | 12446 | 24768 |
| **Longest contig (bp)** | 97957 | 134446 |
| **GC** | 51.59 | 53.41 |

Genome bins quality as accessed by the CheckM tool.

### Symbiotic genome functional annotation and multivariate analysis

Nearly complete genomes gills.bin.1 and gills.bin.4, as well as other related gammaproteobacterial genomes, including representatives of the Cellvibrionaceae family, and more specifically, the *Teredinibacter* clade (Suppl. Table 7) were automatically annotated at the RAST server [35] under default settings. Carbohydrate-active enzymes (CAZymes) were annotated using the dbCAN web server [36]. Results were filtered for E-value of 1e^-5^ and domain coverage fraction > 0.7. Using such parameters, reannotation of *T. turnerae* T7901 genome leads to recover of 99% of the CAZy domains, with false positive and false negative rates of ~8 and ~7% respectively. Natural products Biosynthetic Gene Clusters (BGCs) were annotated using antiSMASH bacterial version [37] with RAST GenBank generated files as input and default settings. The genes composing the resistome (i.e. genes related to antibiotic resistance) were annotated and quantified with RGI (Resistome Genes Identificator) tool of the Comprehensive Antibiotic Resistance Database (CARD) [38], under the *discovery* criteria for detection of perfect, strict and loose hits. We then performed Principal Component Analysis (PCA) to cluster the selected genomes according their diversities of BGCs and resistance genes, and supervised Random Forest (RF) [39] to highlight the fifteen most influential vectors in each case, using ggbiplot R library (Vu, 2011) to plot the results. The number of variables in PCA was reduced by supervised RF to, with higher mean accuracy using the R library [40]. To test if the relative abundance of resistome and metabolome (specifically Nrps, PKS and their hybrid paths) related genes were significantly different between free-living and host-associated bacteria we performed Kruskal-Wallis non-parametric test using the kruskal.test() function in R and considered p-value lower than 0.05 significant. We tested each category independently and adjusted the p-value using the Bonferroni adjust method with the p.adjust()function in R.

### Phylogenetic analysis

The two high quality genome bins were placed in a phylogeny with closely related Gammaproteobacteria based on the 43 phylogenetically informative marker genes that CheckM uses to assess bin placement in its own backbone tree (Supplemental Table S6 in [34]). All genomes from the family Cellvibrionaceae were downloaded from the PATRIC (Pathosystems Resource Integration Center, [41] genome database, to our knowledge the largest publicly available prokaryotic genome database. One genome per family of Halieaceae, Porticoccaceae, and Spongiibacteraceae was added to together serve as outgroups. It was recently shown that the monophyly of Cellvibrionaceae could only be clearly resolved when using a genome-scale supermatrix [42] while a tree based on the RpoB gene showed Cellvibrionaceae to be paraphyletic with Microbulbiferaceae. Thus, no Microbulbiferaceae genomes were included here. Indeed, when Microbulbiferaceae were included in the tree, this led to paraphyly of the Cellvibrionaceae, but the placement of the novel genome bins within the Cellvibrionaceae remained unchanged (not shown). Genomes were downloaded based on the genome_lineage file on the PATRIC server that was downloaded on December 20, 2017. One Cellvibrionaceae genome present in the genome_lineage file, *Cellvibrio mixtus* strain PSBB022, could not be retrieved from the PATRIC server. CheckM was run on all the genomes to identify the 43 marker genes, translated sequences were individually realigned after dealignment with Clustal Omega 1.2.3 [43] and subsequently concatenated, filling gaps if a gene was not found in a genome. A maximum-likelihood tree was inferred with RAxML 8.2.9 [44] using the PROTCATLG model and 100 rapid bootstraps (random seeds -p and -x were both set to 12345), and the best scoring tree was visualized with iToL [45]. Additionally, the partial nucleotide sequences of the protein-coding gene *rpoB* (1,020 bp) and *gryB* (1,053 bp) were retrieved from a previously determined set of 25 *T. turnerae* isolates [8], and aligned with respective gene fragments from binned genomes gills.bin.1 and gills.bin.4 with Molecular Evolutionary Genetics Analysis (MEGA) software version 6.0[46], using the built-in ClustalW program with default settings for “*codons*” (protein-coding sequences). Maximum likelihood fits for 24 different nucleotide models were evaluated for each gene and best-fit model of evolution selected according to the lowest Bayesian Information Criteria (BIC) scores. Neighbor-joining phylogenies were reconstructed employing Kimura 2-parameter nucleotide substitution model improved with 5 discrete gamma categories for a total of 1000 bootstraps.

## Results

### The *N. reynei* gill symbiotic microbiome is dense and highly dominated by -proteobacteria, while digestive glands and intestine present similar diverse microbial communities that do not overlap with the gill community

To extend knowledge regarding the role of shipworm symbiotic bacteria in wood digestion and host defense, we performed functional metagenomics of *N. reynei* digestive gland, intestine, and gill tissues (see methods). Shotgun sequencing of fifteen metagenomes samples (Table S1), generated 21,548,783 reads, from which approximately 92% (3.94 Gb) passed the quality control and annotation pipeline of the MG-RAST server. Approximately 1.2, 55.7 and 3.8% of the quality proofed data presented matching hits for ribosomal RNA, protein, and protein with known functional coding genes respectively (Table S1).

Unsupervised multivariate analysis showed that the *N. reynei* gill microbial community has unique taxonomic and functional profiles (Figure 1). Hierarchical clustering of the microbial annotations formed consistent cladogram topologies, with gills samples grouping apart from digestive gland and intestine samples in two main clades (Figure 1A and 1B). Indeed, gill samples exhibited a dense but low diversity microbiome, with Bacteria dominating the hits against the RefSeq database (> 98%), and the vast majority of these hits being tracked to γ-proteobacteria (> 92%), and ultimately, to the genus *Teredinibacter* (Figure 1A, Figure S1). Accordingly, 16S rRNA annotation against the RDP database confirmed the dominance of signatures of *Teredinibacter* in the gill microbiomes (~ 88%) (Suppl. Fig. 1B). Intestine and digestive gland samples, on the other hand, presented more diverse microbiomes distinct from the gills, responding respectively to 23,8 and 28,7% of the taxonomical annotations (Figure 1A, Figure S1).

**Figure 01.**
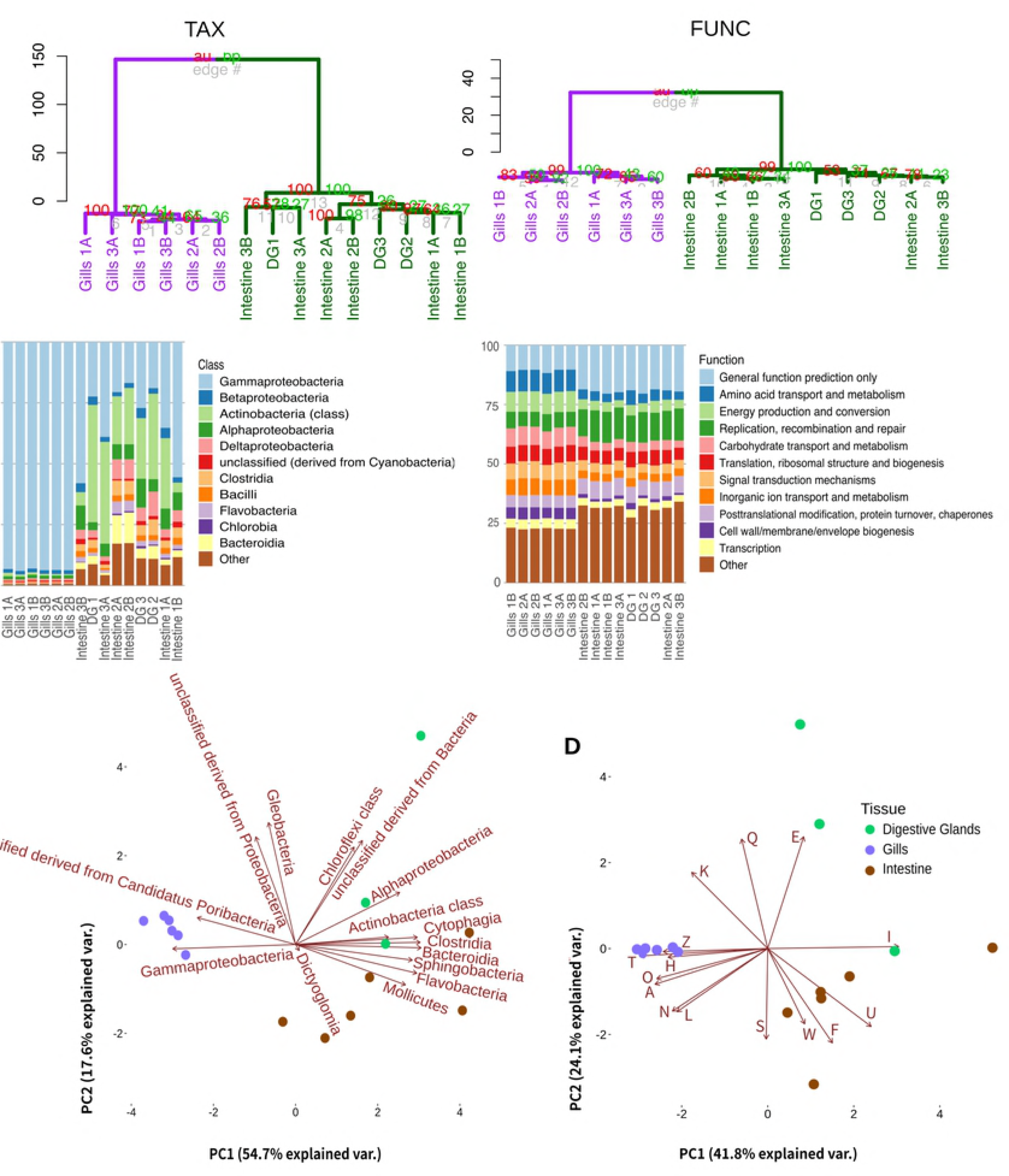
**– Multivariate analysis of metagenomes microbial taxonomical and functional features**. A) Unsupervised hierarchical clustering of taxonomical (TAX) and functional (FUNC) features. Clades formed by gill samples and by intestine plus digestive gland samples are highlighted respectively in purple and dark green. The pvclust AU (Approximately Unbiased) and BP (Bootstrap Probability) p-values are shown at clades nodes in red and light green respectively. B) Relative abundances of metagenome bacterial classes and functions respectively signed with RefSeq and COG (Level 2) databases. C) Multiple dimensional scale (MDS) plot of *N. reynei* metagenomics samples using distances and the fifteen most important bacteria classes (RefSeq database) and functions (COG database at Level 2) vectors as calculated from unsupervised random forest. Digestive gland, intestine and gill samples are shown as green, brown and purple dots respectively. Clusters of Orthologous Groups are classified into functional categories as follow: A – RNA processing and modification; E – Amino acid transport and metabolism; F -Nucleotide transport and metabolism; H – Coenzyme transport and metabolism; I – Lipid transport and metabolism; K – Transcription; L – Replication, recombination and repair; N – Cell motility; O – Post-translational modification, protein turnover, and chaperones; Q – Secondary metabolites biosynthesis, transport, and catabolism; S – Function Unknown; T – Signal transduction mechanisms; U – Intracellular trafficking, secretion, and vesicular transport; W – Extracellular structures; Z – Cytoskeleton

Non-parametric multidimensional scaling (nMDS) combined with random forest biplot (Figure 1C) showed that enrichments to γ-proteobacteria is a major vector driving the observed grouping of gill samples, while gram-positive (Actinobacteria, Flavobacteriia, Clostridia), gram-negative (α-proteobacteria, Cloroflexia, Bacteroidia, Sphingobacteriia), and unclassified Cyanobacteria influence the grouping of digestive glands and/or intestine samples. Additionally, functions under the COG subcategories *Cytoskeleton; Cell motility*; *Coenzyme transport and metabolism*; *RNA processing and modification*; *Signal transduction mechanisms*; *Posttranslational modification, protein turnover, chaperones*; *Transcription*; and *Secondary metabolites biosynthesis, transport and catabolism* influenced gill samples grouping, while digestive glands and intestine microbiomes were driven by *Lipid; Amino acid, and Nucleotide transport and metabolism* besides *Extracellular structures*, and *Intracellular trafficking secretion and vesicular transport* (Figure 1D). The two group comparisons performed with reference to the two main clades obtained by the hierarchical clustering analysis and using taxonomical and functional microbial sequence annotations corroborate the nMDS analysis, with biologically significant enrichments detected in either gills or digestive tract (digestive glands and intestine) microbiomes closely matching the vectors driving samples grouping on nMDS plot (Figure S2).

Although highly informative, comparing taxonomic and functional annotations confine the sequence data to the reference databases, therefore biasing the results according to the database representativeness. Considering that, we further performed annotation-independent comparative metagenomics with crAss [25], a tool that accesses (and score) similarities between metagenomes according to their shared entities (cross-contigs) after cross-assembling all sample’s reads. Interestingly, metagenomes were grouped into one of two different cladograms topologies reconstituted from four distance matrices (Figure 2). When qualitative comparison was applied using “minimum” and “shot” presence/absence-based distance formulas, the samples grouped significantly by the animal source (p<0.001) (Figure 2A). Although, when more quantitative distance formulas “reads” and “wootters” were used, in which the contig number of reads is considered as reflecting its abundance in the environment, samples grouped significantly (p<0.001) by the tissue, driven by gill samples, which clustered together regardless of the animal source (Figure 2B).

**Figure 02.**
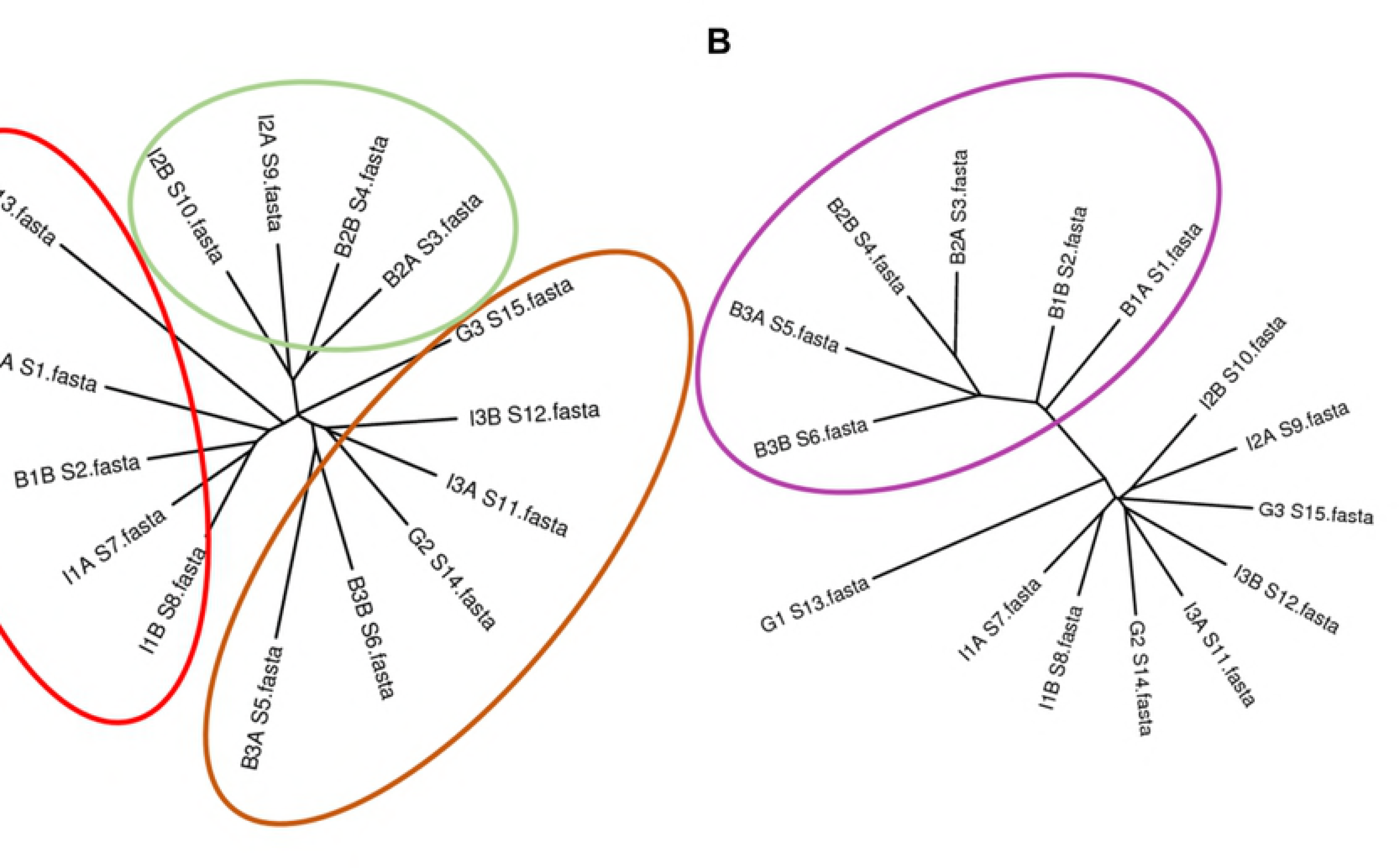
**– Comparison of** ***N. reynei*** **metagenomic samples by cross-assembling.** A) Cladogram representing cross-contigs (i.e., shared contigs containing reads from at least two metagenomes) grouping when using presence-absence-based formulas (distance formulas ‘shot’ and ‘minimum’). Samples grouped according the specimen of origin, being clades formed of samples of *N. reynei* specimens #1, 2 and 3 highlighted in red, green and brown respectively. B) Cladogram representing cross-contigs grouping when using more qualitative distance measures (distance formulas ‘Wootters’ and ‘reads’). Gill samples grouped together regardless the specimen of origin, as highlighted in dark-blue.

Further, we applied the **c**ontigs **a**nnotation **t**ool (CAT) algorithm to get taxonomical classification of contigs assembled for each tissue-type dataset (see Methods). Such analysis showed that gill samples contain 10 times more bacterial-derived contigs (~25% of the signatures) than intestine and digestive glands (~2-3% of the signatures) (Figure S3). Mollusca represented 30 – 38% of the obtained classifications being the most prevailing annotation on the three tissues (Figure S3). Corroborating raw reads annotations, gammaproteobacteria dominated the gill microbial community (~87% of bacterial contigs), with *Teredinibacter* as the most abundant classified bacterial genus (~81% of gammaproteobacterial reads). Additionally, the discrete microbiomes of digestive glands and intestine were reaffirmed as more diverse with Proteobacteria (~0.6 – 0.7%), Actinobacteria (~0.4 – 0.5%), Bacteroidetes (~0.3 – 0.6%) and Firmicutes (~0.3%) contigs leading the obtained microbial hits (Figure S3).

### *T. turnerae* and *Teredinibacter* sp. compose the *N. reynei* gill symbiotic community

Genome binning was then performed in an effort to recovering microbial genomes representing each investigated tissue microbiome. Two γ-proteobacterial genomes bins presenting the quality required for further analysis (> 50% completeness, < 5% contamination), as assessed by CheckM tool [34], were recovered, both from the gill metagenome dataset (Table 1). The genome bins, labeled as “gills.bin.1” and “gills bin.4”, were automated assigned as *T. turnerae* and *Teredinibacter* sp. respectively (Table 1).

Firstly, the taxonomic affiliations of gills.bin.1 and gills.bin.4 genome bins were checked by genome-scaled supermatrix (Figure 3) and one-protein-coding taxonomic marker phylogenies (Figure S4). Corroborating with the CheckM taxonomical annotation, which is based on lineage-specific marker genes, both phylogenies provided highly similar tree topologies where both genome bins fell in a subclade of Cellvibrionaceae family, with gills.bin.1 grouping within the previously defined “clade 1” of *T. turnerae* [8] and gills.bin.4 appearing as outgroup of this species main clade (Figure 3; Figure S4). The detection of a *N. reynei* symbiotic “clade I” *T. turnerae* strain is in agreement with previous affiliation of a *N. reynei T. turnerae* isolate, T8508, also falling within this clade [8]. Moreover, as observed for other Teredinidae species, the present results showed that *N. reynei* gill symbiotic microbiota contains species in the *Teredinibacter* clade including, but not limited to, *T. turnerae*.

**Figure 03.**
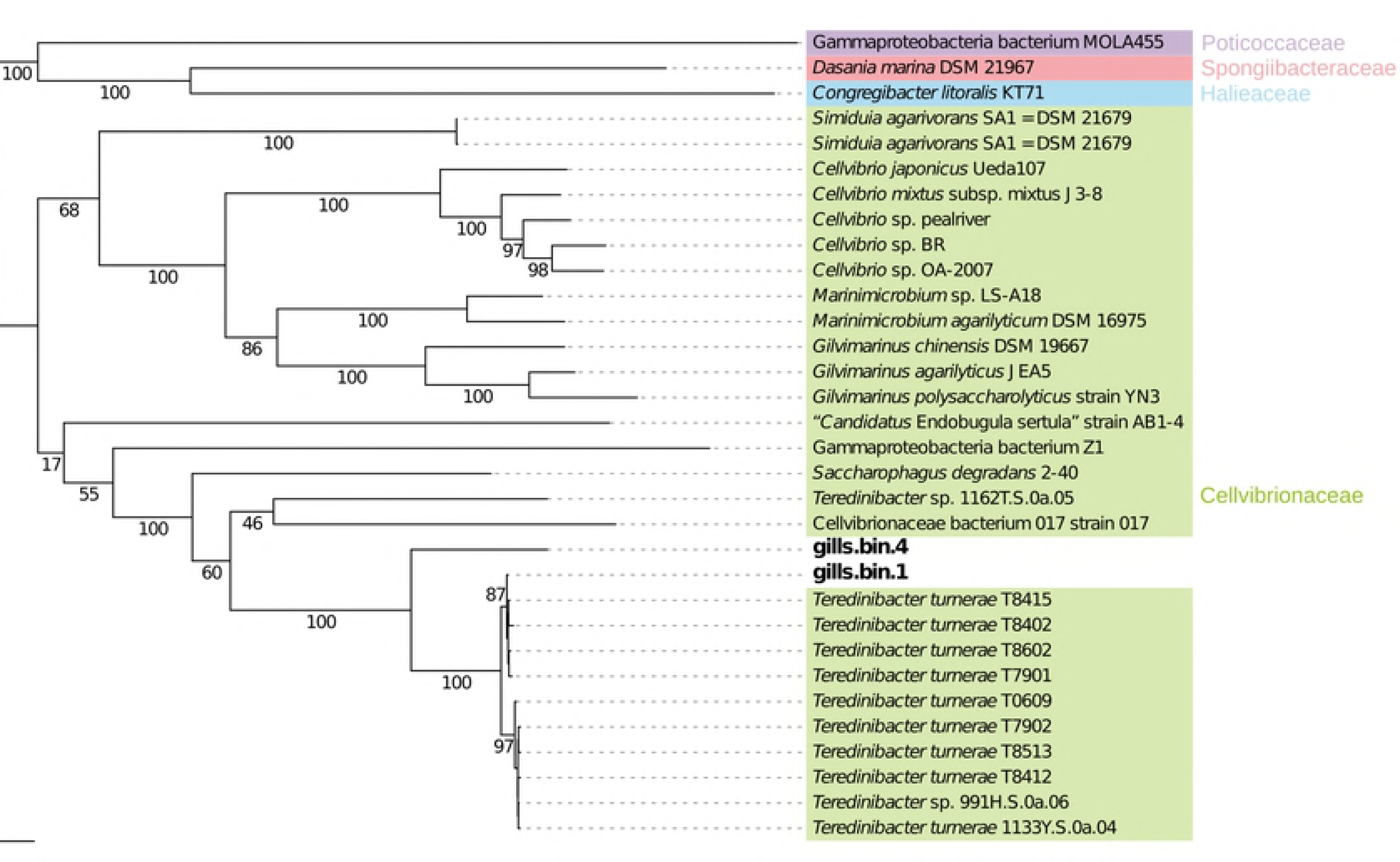
**– Phylogeny of two high quality Teredinibacter bins based on a concatenated alignment of 43 proteins.** RAxML maximum likelihood tree based on the CAT model of rate heterogeneity and LG amino acid substitution matrix, with 100 rapid bootstraps. To improve readability, bootstraps for the *Teredinibacter turnerae* sub-clades I and II (with bootstrap values of 87 and 97 respectively) are omitted. The tree is rooted in between the Cellvibrionaceae branch and representatives of Halieaceae, Porticoccaceae, and Spongiibacteraceae. Color coding of the families is similar to the one used by Spring and co-authors (2015). The scale bar represents 0.05 nucleotide substitutions per site.

### *N. reynei* symbiotic *Teredinibacter* genome bins code for wood specialized repertoire of hydrolases including novel multicatalytic enzymes

The *T. turnerae* T7901 genome codifies an arsenal of enzymes devoted to breaking down the lignocellulosic plant material [9]. Based on this knowledge, we annotated the recovered genome bins against the carbohydrate-acting enzymes (CAZy) database [47] and compared their cazymes profile with the one for T7901, disposed at the CAZy server (Table S2). In the three compared genomes, cazymes represented between 3.5 – 4% of the protein coding genes, encompassing a diversity of predicted protein domains including 32 – 37% of glycoside hydrolases (GH), 9 – 12% of glycoside transferases (GT), 7 – 13% of carbohydrate esterases (CE), 1 – 2% of polysaccharide lyases (PL), and 42 to 44% of carbohydrate binding modules (CBM) (Table S2). Additionally, when GH families were grouped according their substrate specificities, it was shown that the binned genomes also presented the “wood-specialized” profile described for T7901 [9], where a major portion (43 – 51%) of the GHs are devoted to the digestion of wood-composing polysaccharides (Figure 4A). Furthermore, gills.bin.1 contains identical or near identical (E value = 0.0, amino acids identities ≥ 99%) homologs to the multidomain and multi-catalytic cazymes detected in the T7901 genome, the exception being TERTU_RS21420, coding for the biochemically characterized multifunctional cellulase CelAB [48] (Table S3). In comparison, gills.bin.4 codes less conserved homologs (E value = 0.0, amino acids identities ≥ 75%) to only three of T7901 multicatalytic enzymes (Table S3). However, this genome bin codes for other novel multidomain/multicatalytic enzyme configurations, including a putative acetyl xylan esterase/ endo-1,4-β-xylanase (peg.397) and two putative dual pectin acetylesterases (peg.2052 and peg.3260) (Figure 4B).

**Figure 04.**
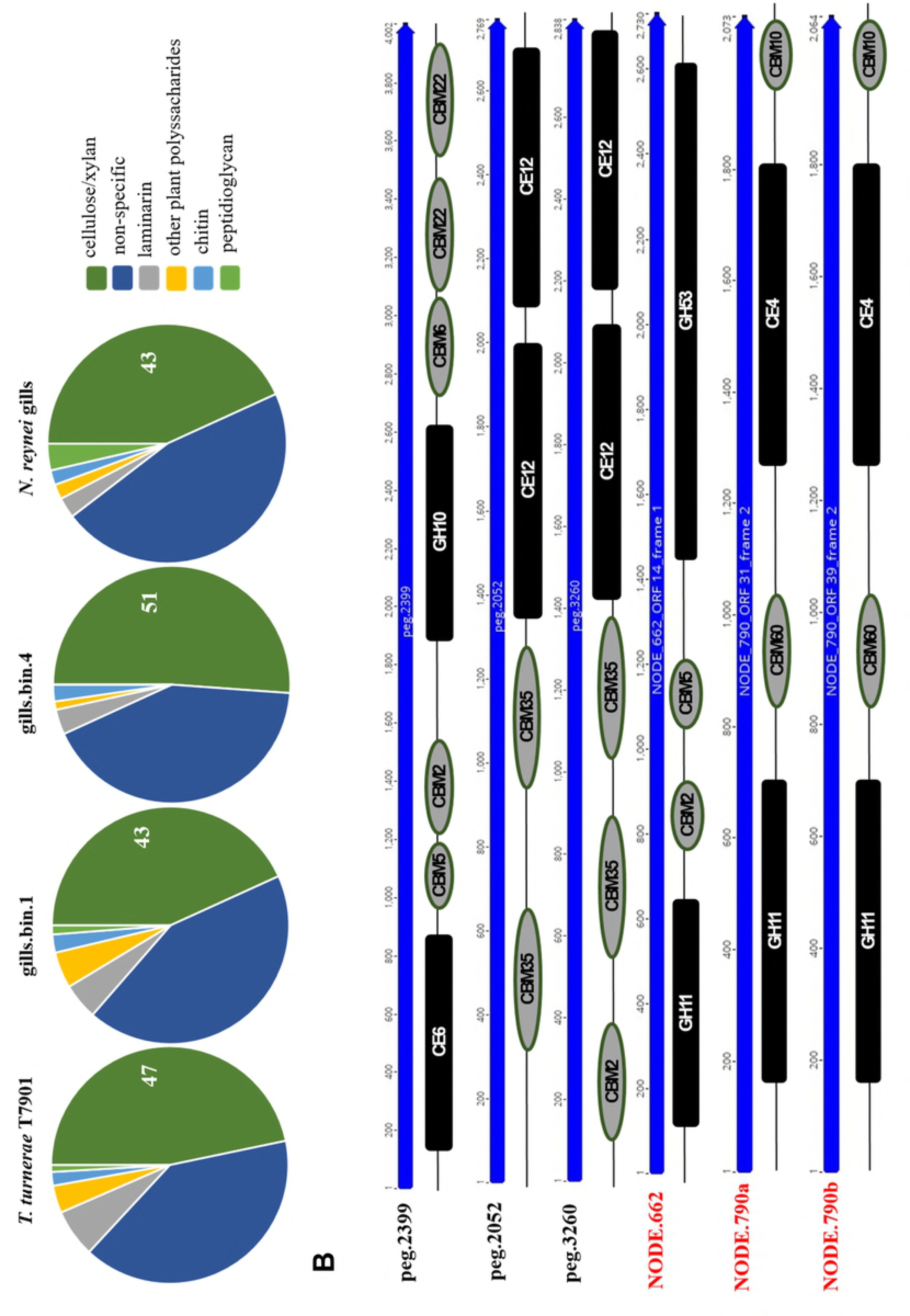
**– Gills genome bins and unbinned contigs carbohydrate active enzymes (CAZymes).** A) Proportion of GH domains according to the substrate specificity at gills.bin1 and gills.bin 4 binned genomes and at other unbinned contigs from gills metagenome dataset. Substrate specificities as previously defined [9]: dark green = cellulose/xylan GH families (5, 6, 8, 9, 10, 11, 12, 44, 45, 51, 52, 62, and 74); dark blue = GH families with other or not-unique specificities (1, 2, 3, 13, 15, 23, 27, 30, 31, 35, 39, 43, 73, 77, 78, 79, 94, 95, 103, 108, 109, 115, 128, 130); light grey = laminarin GH families (16 and 81); yellow = other plant cell wall polysaccharides GH families (26, 53, and 67); light blue = chitin GH families (18, 19, and 20); light green = peptidoglycan GH families (28 and 105). B) Novel multicatalytic CAZymes configurations detected on *Teredinibacter sp.* gills.bin.4 genome bin (labeled in black), and on unbinned contigs from gill-symbiotic gammaproteobacterial community (labeled in red). Abbreviations key: glycoside hydrolases (GH), carbohydrate esterases (CE), carbohydrate binding modules (CBM). Catalytic domains and binding modules are color coded as black squares and gray circles.

When the entire set of contigs assembled from the gill dataset was mined, additional 291 putative cazymes not mapped to the genome bins were found. In total, *N. reynei* gills contained 882 carbohydrate active domains, and comparing to the cazymes profile of the symbiotic genome bins, the gill dataset presented an ~2.3 X enrichment for GTs, and 1.3 X reduction on GHs (Suppl. Table 2). However, the wood-specialized profile is maintained for the entire gill symbiotic community CAZymes (Fig. 4A). In fact, other unique multicatalytic woody plant material digesting CAZymes that have not binned to *Teredinibacter* genomes were detected, such as a putative endo-β-1,4-xylanase/ endo-β-1,4-galactanase (NODE.622) and two endo-β-1,4-xylanase/acetylxylan esterases (NODE.790a and b) (Fig. 4B).

Finally, we evaluate genomes CAZymes in light of the recent genome-enabled-proteomic study performed with shipworm specimens of *B. setacea* species, which revealed that an array of carbohydrate-active-domains derived from the gill symbiotic gammaproteobacterial community are found in the animal’s cecum contents [11]. We verified that, in the compared genomes and in the gill microbiome dataset, the related domains families are enriched, representing 82 – 89% of the cellulase/xylanases (GH families 5, 6, 9,10, 11, 45 and 53), 50 – 75% of the carbohydrate esterases (CE families 1, 3, 4, 6 and 15) and 44 – 62% of the carbohydrate binding motifs detected (CBM2, 10, 22, 57, 60, 61) (data not shown).

Contrasting with the rich and wood devoted repertoire of hydrolyses present in the gill symbiotic site, contigs from digestive glands and intestine datasets retained only 29 and 101 CAZymes related domains respectively, where GHs are absent in recovered enzymes of digestive glands and account for only 4% of intestine-proteins CAZymes, all of which have substrate specificity other than cellulose/xylan (Table S2, data not shown).

### The *N. reynei* symbiotic genome bins code for a diverse repertoire of biosynthetic gene clusters (BGCs) including *Teredinibacter* related and novel putative pathways

Further, we evaluated the potential of the *N. reynei* associated microbiome to produce bioactive secondary metabolites. Firstly, the gill, digestive gland and intestine assembled sequence dataset was mined against the robust shell of algorithms of the antiSMASH server [37]. The gill symbiotic microbiome was shown to be a hotspot of BGCs, with a total of 119 contigs encoding enzymes involved in biosynthesis of complex peptides, polyketides and hybrid compounds, besides poly-unsaturated fatty acids (PUFAs), terpenes, arylpolyenes, homoserine lactones, ectoine and bacteriocins (Figure 5A). On the contrary, no secondary metabolite clusters were detected in the digestive gland dataset, and only two contigs for modular type I PKS were found in intestine samples (data not shown). Intriguingly, the intestine-derived type I PKS contigs best BLASTp hits were to a fusarin C synthetase-like (XP_021367770) and a erythronolide synthase (OWF54747.1) genes linked to the genome of the bivalve *Mizuhopecten yessoensis* (Yesso scallop) [49]. Since modular type I PKS occur almost exclusively in microbes, it is unclear if these homolog genes are truly encoded in the *M. yessoensis* genome, which would indicate a rare inter-kingdom horizontal gene transfer event, or are from a sequenced contaminating DNA source, as an associated bacterium.

**Figure 05.**
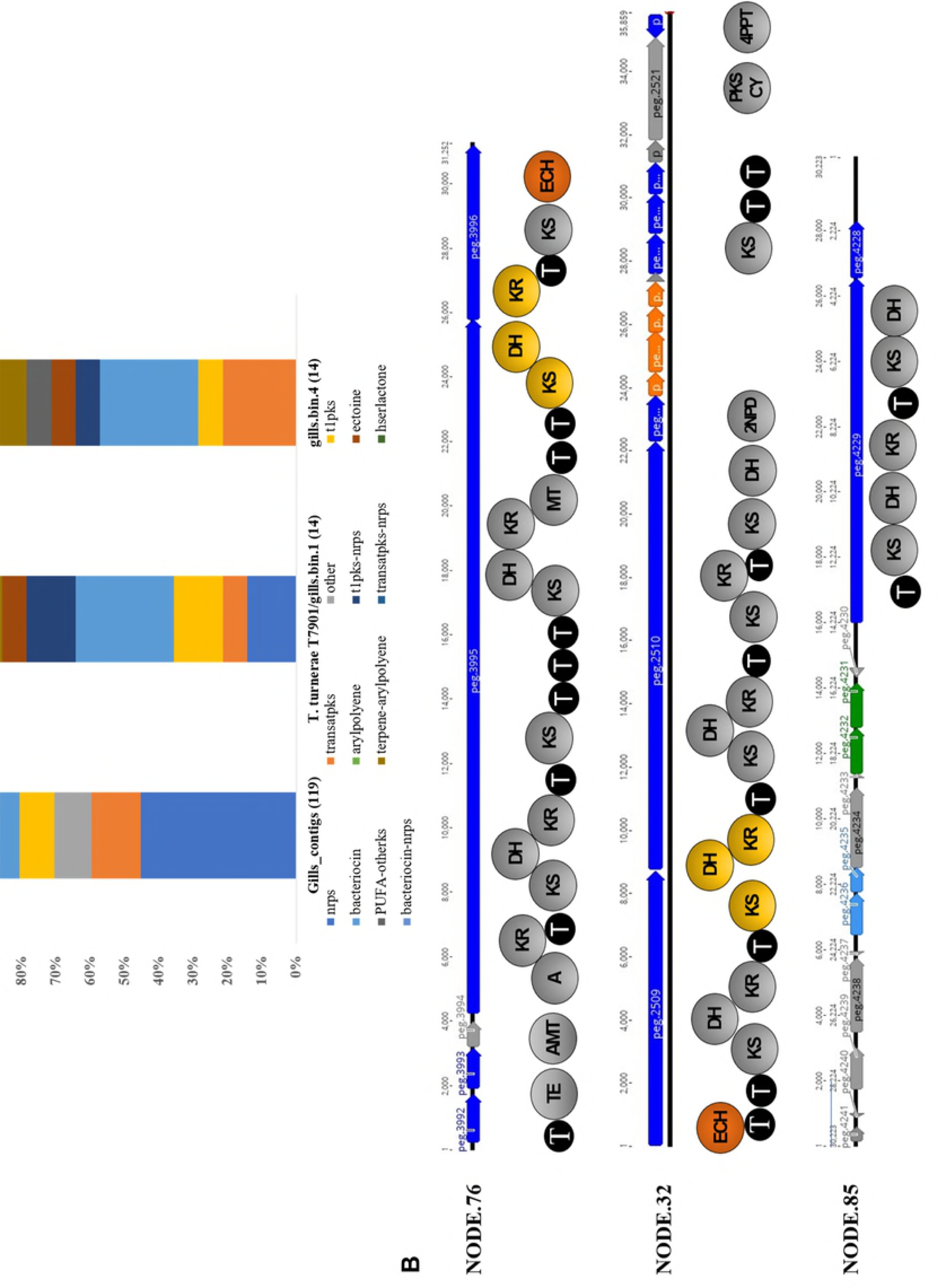
**– Gills genome bins Biosynthetic Gene Clusters (BGCs).** A) Secondary metabolomes. BGCs/contigs detected in *Teredinibacter turnerae* (T7901 and gills.bin.1), *Teredinibacter sp.*(gills.bin.4) and on contigs derived from *N. reynei* gill assembled dataset. Clusters are grouped, and color coded according to the compound class. B) Novel putative *trans-*AT PKS biosynthetic gene clusters detected on *Teredinibacter sp.* gills.bin.4 genome and the main predicted catalytic domains of their open reading frames (ORFs). Biosynthetic, transport-related, regulatory, other β-branching-related genes are color coded in dark-blue, light-blue, green, gray and orange respectively. Domains keys: thioesterase (TE), aminotransferase (AMT), adenylation (A), ketosynthase (KS), ketoreductase (KR), dehydrogenase (DH), thiolation (T), enoyl-CoA dehydratases (ECH), 2-Nitropropane dioxygenase (2NPD), polyketide synthase cyclase (PKS CY), 4’-phosphopantetheinyl transferase (4PPT). Putative inter-proteins bimodules are present in yellow, PKS-encoded ECH catalytic domains are present in orange.

We then reanalyzed the secondary metabolome of the reference *T. turnerae* T7901 genome via the antiSMASH server. Previously, T7901 was shown to devote an unexpectedly high percentage (~7%) of its genetic content to natural product biosynthetic pathways [9]. The new genome mining reveals a total of 13 BGCs, adding new putative pathways for bacteriocins (4), terpene (1) and ectoine (1), and condensing firstly delimitated “regions” 3, 4, and 5 [9], into |a putative giant Bacteriocin-T1pks-Nrps BGC of ~270.6 kb (Table S4). Screening of gills.bin.1 and gills.bin.4 genome bins lead, respectively, to the detection of 19 and 14 contigs, representing putative BGCs, or fragments thereof, for production of compounds from a diversity of classes such as polyketides, non-ribosomal peptides, terpenes, and bacteriocins (Figure 5A). To get this analysis at higher perspective, we also annotated the secondary metabolome of all genomes in the Cellvibrionaceae family included in our genome-wide phylogeny (Figure 3) and resolved the detected BGCs diversity by supervisioned random forest combined with multidimensional scaling. The *N. reynei* gill symbiotic genome bins contain secondary metabolomes highly related to the ones from representative *T. turnerae* strains, which grouped tightly mostly because of enrichments for modular PKS, NRPS and hybrid pathways (Figure S5).

Eighteen of the putative BGCs from gills.bin.1 and additional fourth contig undetected by the antiSMASH server could be mapped to all 13 BGCs from *T. turnerae* T7901 secondary metabolome with high pairwise identity (~99%) and BGCs coverage (~88%) (Table S4). BLASTp inspection showed that the type 1 PKS route from gills.bin.1 genome bin lacking in T7901 genome is actually conserved in other *T. turnerae* genomes, as the ones from strains T8412, T8413 and T8415 (E value = 0, 100% query coverage, ~99% sequence identity) (Table S5). Therefore, such results corroborate the grouping of gills.1 genome bin as a *T. turnerae* strain.

Conversely, only a putative bacteriocin BGC encoded in gills.bin.4 could be mapped to the *T. turnerae* T7901 secondary metabolome (Suppl. Table 4). BLASTp analysis showed that eight others putative BGCs from this *Teredinibacter* sp. genome bin were also conserved (~97% gene coverage and ~73% identity) in other *Teredinibacter* genomes (Table S5). The 5 remaining contigs from gills.bin.4 are novel among the secondary metabolomes of shipworms symbionts.

The core biosynthetic proteins encoded on these BGCs presented best BLASTp hits to putative proteins from Bacteroidetes (~ 40%), gammaproteobacteria (33%), and betaproteobacteria (~15%), besides alphaproteobacteria and Firmicutes (Table S6). Three of these hypothetical novel BGCs (NODE_76, NODE_26 and NODE_85) code for typical multi-modular *trans*-AT PKS enzymes, with a catalytic domain inventory including two inter-proteins bimodules, commonly involved in formation of α,β-olefinic moieties, and a gene cassette typically involved in the formation of β-branching in bioactive compounds such as the highly cytotoxic pederins and the antibiotic bacillaene [50] (Figure 5B).

### The *Teredinibacter* resistome singularity is driven by putative resistance genes to PKs-NRPs derived antibiotics

The Resistome is defined as the collection of all antibiotic resistance genes present in a microorganism’s genome, including precursor genes, which encode proteins retaining weak antibiotic resistance or binding activity, and that under selective pressure can evolve into a new resistance marker [51]. Here we also investigated the resistome coded in *Teredinibacter* genomes, including gills.bin.1 and gills.bin.4 genome bins, and other representative genomes of the Cellvibrionaceae family. The vast majority of resistance signatures obtained with the Resistance Gene Identifier (RGI) tool from the Comprehensive Antibiotics Resistance Database (CARD) [52] were under the lower stringency cutoffs, which includes putative resistance genes precursors (Table S7). Multivariate comparisons showed that *Teredinibacter* resistomes grouped tightly and are particularly driven by putative resistance markers to fluoroquinolone (total of 2110 hits) and to antibiotics derived from PKS, NRPS and hybrid compounds, including macrolides (2071), monobactam (2071), tetracycline (1963), penam (1343), peptides (816) and glycopeptide (739) antibiotics (Figure 6A). The relative abundance of genes encoding resistance related proteins was not significantly different (adjusted p-value of 0.28) across the genera inside the Cellvibrionaceae family, suggesting that the resistome size is conserved (Fig 6B, yellow bars). Indeed, resistomes of both *Teredinibacter* (13) and *Cellvibrio* (6) representative genomes were of similar size (representing respectively ~5.3 and ~5.6% of the protein coding genes) (Table S7), despite the intracellularly symbiotic life-style of the former and, mostly free-living nature of the latter. In contrast, BGCs and, more specifically, Nrps-PKS-hybrid metabolic paths relative abundances differed significantly between host-associated and free-living Cellvibrionaceae (adjusted p-values <0.001 and <0.0001, respectively), being larger in the first group (Figure 6B, and 6C). Clearly this pattern is driven by symbiotic *Teredinibacter* and, mostly free-living *Cellvibrio* representative genomes (Figure 6B).

**Figure 06.**
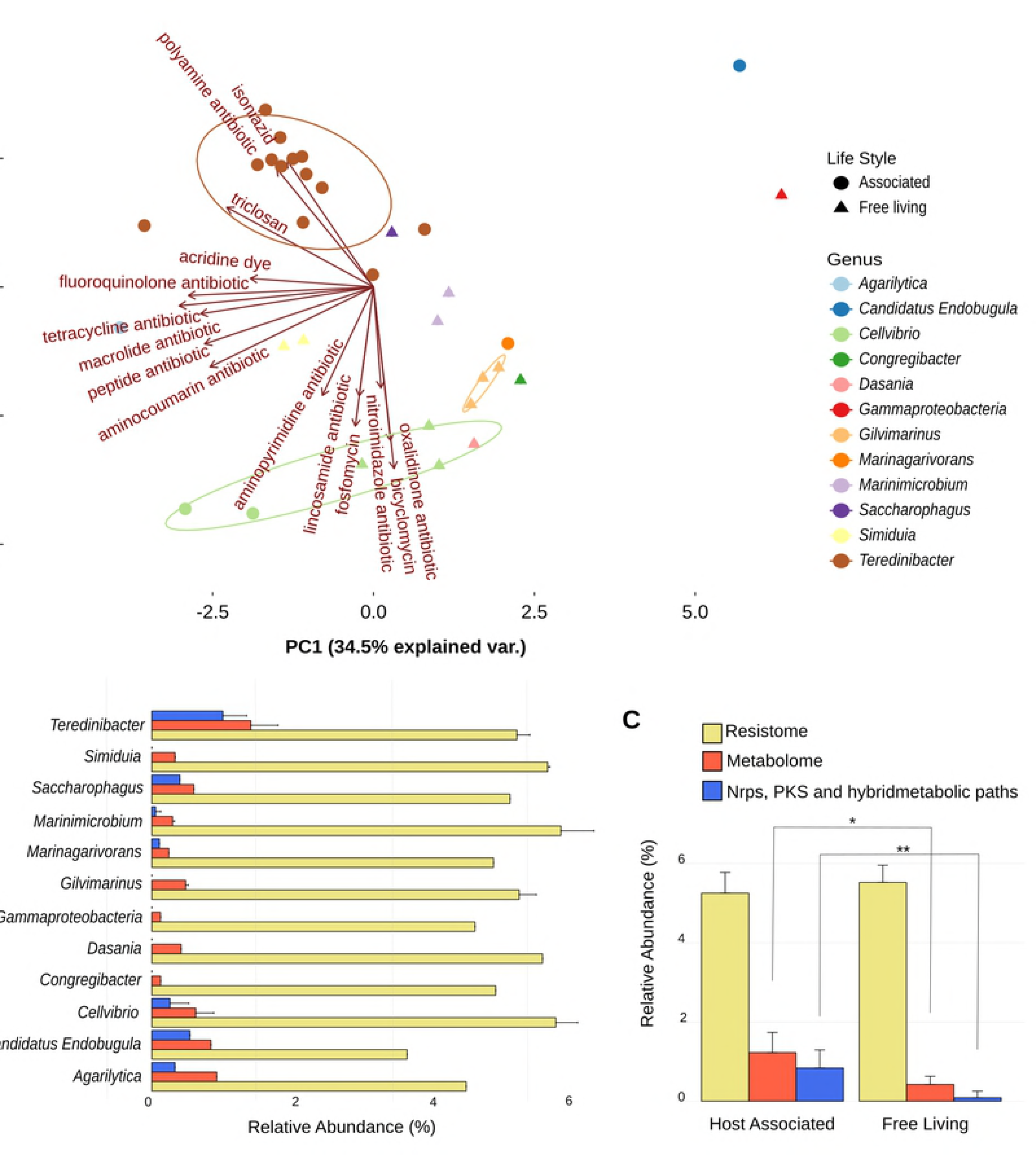
**–Teredinibacter resistome.** A) resistome PCA using the fifteen most important genes based on mean decrease accuracy of Random Forest analysis. *Teredinibacter* forms a major group influenced by polyamine, isoniazid, and PK-NRP-derived antibiotics. B) Relative abundance of resistance gene markers, BGCs, and NRPS, PKS and hybrid BGCs. C) Host associated bacteria present a significantly larger metabolome and abundance of NRPS, PKS and their hybrid metabolic paths than free living bacteria. One asterisk represents Kruskal-Wallis test p-values <0.001 and two represent <0.0001 both adjusted with Bonferroni method.

## Discussion

In the present work, metagenomics was applied to characterize, compare and investigate the microbiomes associated with the digestive glands, intestines and gills of the mangrove shipworm *N. reynei*. Taxonomic annotations showed that, as seen for other shipworm systems [3,4,11], the gills of *N. reynei* are a unique symbiotic site, filled with cellulolytic γ-proteobacteria, including bacterial types of the *Teredinibacter* clade (Figure 1, Figure S1 and S3).

In contrast, digestive gland and intestine samples differ from the gill community and are more diverse (Figure 1, Figure S1 and S3). Such results corroborate previous culture-independent analysis of five shipworms species microbiomes, where digestive tract tissues were reported as virtually sterile (cecum) or containing very low loads of bacteria (intestine) [15].

Therefore, our results indicate that digestive tract microbiomes do not contribute significantly to the digestion of wood particles and consequently the nutrition of the shipworm host. Remarkably, shipworms that are basal in the Teredininae subfamily (including *N. reynei*, *Dicyathifer manni* and *Bactronophorus thoracites*) have the anus opening into the anal canal structure [1,2], which morphology indicates a role in animal’s food absorption [18], but microbial constitution is yet to be explored.

*N. reynei* gill samples are enriched for functions under categories of cellular process and signaling (e.g., *Cell motility*; *Signal transduction mechanisms*), information storage and processing (e.g., *RNA processing and modification*, *Transcription*) and metabolism (e.g., *Inorganic ion transport and metabolism*, *Iron acquisition and metabolism*, *Nitrogen metabolism*, and *Secondary metabolites biosynthesis, transport, and catabolism*) (Figure 1D, Figure S2).

Cross-assemblage based comparative metagenomics showed that such enrichments were related to gammaproteobacterial-derived genes which drive gill samples to group together regardless of the shipworm species of origin (Figure 2B). In fact, gill-enriched functionalities are consistent with the genetic repertoire described for the prototypical shipworm symbiont *T. turnerae,* a flagellar gammaproteobacterial species able to fix dinitrogen, sequestering iron and producing secondary metabolites *in symbio* [9,10,12].

Specific mining of the contigs assembled from each tissue dataset showed that the *N. reynei* gill microbiome is a hot spot of genes for: i) CAZymes, and particularly, wood-degrading hydrolases (Figure 4), and ii) biosynthetic pathways for a variety of bioactive compounds classes (Figure 5). On the other hand, such a genetic repertoire was shown to be negligible in digestive gland and intestine datasets. However, is important to note that the digestive tract microbiomes have proven to be more diverse than the gill symbiotic gammaproteobacterial community, and so, our sequence strategy loses resolution and may not reach the necessary sequencing depth required to fully cover these microbes’ functionalities.

Two high-quality representative symbiotic genomes could be recovered from the gill datasets by our binning strategies (Table 1), and were assigned as *T. turnerae* and *Teredinibacter* sp. by CheckM tool sets of single-copy genes that are signatures within a phylogenetic lineage [34]. A phylogeny with several closely related reference taxa confirmed these taxonomical affiliations grouping gills.bin.1 and gills.bin.4 genome bins respectively inside and as a sister group of the *T. turnerae* species clade (Figure 3). The tree topology reliably reproduced the one originally obtained by Spring and co-authors (2015), which supported the proposal of new gammaproteobacterial families, including the Cellvibrionaceae family embracing *Teredinibacter* and *Ca.* Endobugula genera of marine invertebrate symbionts [42].

Detection of *T. turnerae* and *Teredinibacter* sp. as prevalent bacterial types within *N. reynei* gill microbiome is in agreement with previous culture efforts made with this species specimens from the very same sampling site, and that resulted in isolation of several *T. turnerae* strains [19]. These results also agree with previous culture-independent approaches applied on *L. pedicellatus* and *B. setacea* shipworm species, showing that gill gammaproteobacterial communities were composed of at least four distinct bacterial types, all grouping with *T. turnerae* in a main clade of shipworm symbionts [3,4,14].

We also reproduced a systematic phylogenetic analysis previously performed with *T. turnerae* isolates from a wide variety of shipworms species and using protein coding gene markers [8] (Figure S4). In this work, Altamia and collaborators showed that *T. turnerae* population actually encompasses two distinct lineages that, although they experienced horizontal gene transfer events, can be consistently grouped in two main clades: clade I and clade II. Our phylogeny reconstitutions using *rpoB* and *gyrB* markers replicate this topology, also agreeing with the genome-wide phylogeny. In both analyses, gills.bin.1 genome bin fell within *T. turnerae* “clade I”, just as the *N. reynei* isolated *T. turnerae* strain T8508 [8,19] (Figure 3, Figure S4).

Specific mining revealed that *N. reynei*’s symbiotic *Teredinibacter* genome bins contain a repertoire of carbohydrate-active enzymes (CAZymes) that is closely related to the “wood-specialized” profile described for *T. turnerae* T7901 [9], and is characterized by a vast repertoire of cellulases/xylanases and other catalytic and binding domains involved in a network for breaking down woody complex polysaccharides (Figure 4). Of particular interest was the detection of multi-catalytic CAZymes with novel domain configurations in gills.bin.4 and unassembled gammaproteobacterial contigs (Figure 4B). Also, within the cabal of CAZymes characterized was an enrichment of catalytic domains that, in the *B. setacea*, were proven to travel from the gill symbiotic site up to the animal cecum, where the wood digestion takes place [11]. Considering that reduced gills, and well-developed cecum and anal canal organs, both associated with the digestion of wood particles, support the hypothesis that *N. reynei* nutritional depends heavily on wood [17], a symbiotic role on shipping key hydrolases from the gills site to the cecum “biodigester” is quite reasonable and very exciting. Therefore, additional efforts focusing on isolation and discovery of new symbiotic bacterial types from *N. reynei* gills is worthwhile and can lead to the discovery of new and biotechnologically relevant wood-digesting CAZymes.

*N. reynei* gill’s symbiotic microbiome was also shown to be a fruitful source of BGCs. Particularly, the *T. turnerae* representative genome bin (gills.bin.1) was shown to contain all the BGCs previously described for *T. turnerae* strain T7901 [9], including the characterized routes for production of the turnerbactin triscatecholate siderophore [13] and for the tartrolon family of antibiotics [12]. Indeed, these results corroborate the antimicrobial activity previously reported for *T. turnerae* strain CS30, isolated from *N. reynei* [19], and the results of a PCR survey for the tartrolon producing *trans*-AT PKSs (*trtDEF*) that returned expected amplicons when CS30 and other *T. turnerae* isolates genomic DNA were used as template [12]. Additionally, the *in symbio* expression of the *trt* pathway was also shown by reverse transcriptase PCR with *N. reynei* gill bulk RNA (unpublished data). However, the capacity of *N. reynei* symbiotic *T. turnerae* to produce tartrolons remains to be clarified, since compounds of this family could not be detected on a survey including CS30 strain’s culture chemical extracts [12]. Such clarification is of great interest, since it was suggested that tartrolon antibiotics could play a role on shipworm host chemical defense, and/or gill symbiotic community structuring. Finally, growths of CS30 under iron starvation leads to secretion of siderophores, as revealed by the universal chrome azurol S (CAS) assay (unpublished data). Such results support also the production of turnerbactin by this *T. turnerae* strain.

Further, the genome bin gills.bin.4 encoded five contigs representing novel clusters among shipworm microbiomes secondary metabolomes (Figure 4, Table S5). BLASTp searches showed that these clusters putative core biosynthetic proteins present higher similarities to proteins from putative BGCs from Bacteroidetes (as Flavobacteria), betaproteobacteria (as Nesseriales) and firmicutes (as Bacillus) (Table S5 and data not shown). Even knowing that blast searches are biased by the database, and that the modular enzymology commonly involved on biosynthesis of secondary metabolites, such as polyketides, contain many conserved catalytic domains and so, are not good taxonomic markers, these results may be taken as a suggestion of horizontal gene transfer (HGT) events that led to the acquisition of such pathways by the *Teredinibacter* symbiotic community.

Independently of the acquisition history, the discovery of these novel BGCs reinforces the potential of *N. reynei*’s symbiotic system for biotechnological exploration aiming for discovery of new drug leads. In fact, our comparative analyses with representative genomes composing the Cellvibrionaceae family showed that shipworm gammaproteobacterial symbionts, and *Teredinibacter* in particular, share large secondary metabolomes, enriched for PKS-NRPS routes (Figure 6B, Table S7). Such a result reinforces the hypothesis that production of bioactive metabolites, as polyketides and complex peptides or nonribosomally derived siderophores increase *Teredinibacter* fitness *in symbio*. Furthermore, considering that each shipworm species has a singular morpho-physiology and each symbiotic gammaproteobacterial community has a unique diversity, specific BGCs might play specific roles within each host-bacteria interaction.

Consequently, the *trans*-AT PSK enzymology detected in the *Teredinibacter* sp. genome bin is of great interest, since its includes a cassette containing catalytic activities for formation of β-branching chemical structures, which is also present in important drug leads as bryostatins -potent protein kinases modulators, the pederin family of antitumor compounds and myxovirescin antibiotics [50]. Interestingly, macrolide lactones of the bryostatin family, with promising applications for treatment of Alzheimer’s disease, had their biosynthesis assigned to a *trans*-AT PKSs cluster (*bry*) from *Ca*. Endobugula sertula, an as yet not cultivated symbiont of the bryozoan *Bugula neritina* [53], which also groups within the proposed Cellvibrionaceae family [42]. Therefore, efforts towards a better characterization of such putative novel trans-AT-PKS systems are encouraged and can go from attempts to isolate strains of *N. reynei*’s symbiotic *Teredinibacter* sp. carrying these clusters, genome sequencing and retrobiosynthetic compound(s) chemical structure prediction, passing through PCR/probe-hybridization screening of such pathways from *N. reynei* gill metagenomic DNA clone libraries, up to attempts to heterologously express this pathway in a domesticated bacterium or engineered *T. turnerae* strain. Certainly, other interesting BGCs could be retrieved from the 119 putative BGCs or gene clusters fragments recovered from *N. reynei* gill dataset.

The detailed characterization of a BGC is a complex task and usually is made retrobiosynthetically, based on the chemical structure of the cluster-associated purified compound, therefore, beyond the scope of this work. However, the obtained results reaffirm shipwormgammaproteobacterial symbiotic interactions as a prolific source for detection of interesting bioactive compounds, including antibiotics. Indeed, our group and collaborators are working on a deeper and full characterization of the BGC catalog of shipworm γ-proteobacteria symbiotic communities, including a pool of DNA sequence from isolates genome DNA and gill metagenome samples from shipworms from several species and geographical locations.

The representative *Teredinibacter* genomes also encoded more putative resistance markers for PKS and NRPS derived antibiotics (Figure 6A). The matching of this hypothetical resistome with the profile of *Teredinibacter* BGCs indicate that at least some of these putative markers might truly be resistance to self-produced antibiotics, and so, further characterizations of these genes are of great interest.

## Conclusion

In this study we applied metagenomics to characterize the microbial communities associated with key organs of shipworm digestive systems, using the mangrove-specialized species *N. reynei* as a research subject. We provided strong evidences that the *N. reynei* gill symbiont gammaproteobacterial community is a hot spot for biotechnologically relevant enzymes, such as hydrolases involved on breaking down wood-derived complex polysaccharides, and biosynthetic gene clusters for production of potentially bioactive secondary metabolites such as complex polyketides, peptides and lipopeptides. Further, the discrete bacterial communities forming the microbiome of the digestive tract tissues seems to have little or no involvement in host nutritional support or chemical defense. Digestive gland and intestine datasets presented a simple repertoire of CAZymes, lacking glycosyl hydrolases for digestion of woody material, and only one fragmented BGC, from intestine samples, with hits to a *locus* of the Yesso scallop bivalve mollusk genome. Two representative symbiotic genomes were recovered from gill symbiotic community and taxonomically assigned as a *T. turnerae* strain and a yet to be cultivated *Teredinibacter* sp. bacterium. The latter encoded novel multi-catalytic CAZymes, and a *trans*-AT PKS BGC containing the catalytic domains implicated in β-branching in related polyketides. Finally, we show strong evidence that *Teredinibacter* is enriched in secondary metabolite pathways when compared to relatives of the Cellvibrionaceae family particularly by an excess of BGCs for complex polyketides and nonribosomal peptides, reinforcing the possible role of these compounds on supporting the symbiosis.

## Acknowledgments

Metagenomic sequencing has been performed at the Genomics and Bioinformatics facility (CeGenBio) of the Drug Research and Development Center (NPDM) of the Federal University of Ceara. Sampling and Genetic Resources Assessment were performed under license IBAMA/MMA/SISBIO numbers 48388-1 and 48388-2. Authors thank the laboratories of microbiology and of microscopy from the National Institute of Metrology, Quality and Technology (http://www.inmetro.gov.br) at Xerém, RJ, Brazil for the support on preliminary samples processing. We also thank professor Dr. Margo Haygood for the critical reading and revision of this manuscript.

## Supporting Information

**Figure S1 – Metagenomes taxonomic diversity.** A) Relative abundance of metagenomes taxonomical signatures under Domain hierarchical level (RefSeq database). B) Relative abundances of metagenomes bacterial genera when considering 16S rRNA reads (RDP database). C) Box-plot of metagenomes Shannon–Weaver index according to the tissue source. Digestive glands (light-blue), and Intestine (green) samples present higher Shannon diversity number, evenness and entropy when compared with Gills (dark-blue) samples. Annotations were performed under MG-RAST server under default stringency parameters (e-value 1e-5, % of identity = 60%, minimal length of 15 and minimal abundance of 1, considering the representative hit).

**Figure S2 – Two groups comparisons between gills and digestive-tract (digestive glands + intestine) metagenomes annotations.** A) Extended error bar plot showing bacterial classes enriched at gills (blue) or digestive tract (orange) metagenomic groups. B) and C) Extended error bar plot showing gills (blue) or digestive tract (orange) enriched bacterial functions when using Cluster of Orthologous (COG) database (B); or Subsystem Technology database (C), at their respective hierarchical level 2. Two groups comparisons were performed at the STAMP software version v2.1.3, as explained at *Methods*.

**Figure S3 – Contigs based taxonomical annotation of** ***N. reynei*** **gills, digestive glands and intestine datasets.** A) Pizza chart showing relative abundances of taxonomical hits per superkingdom. B) Distribution of hits to the most abundant taxon under the phyla taxonomical hierarchy. Inset on gills bar chart brings the distribution of proteobacterial-derived hits (orange bar) as a pizza chart to show the prevalence (98,2%) of hits to the γ-proteobacteria class.

**Figure S4 – Neighbor-joining phylogenies inferred for the genome bins protein coding gene markers** ***gyrB*** **(A) and** *rpoB* **(B) against a referential dataset of genes from** ***T. turnerae*** **isolates.** Bootstrap probability values > 0.7 are shown at the nodes.

**Figure S5 -Secondary metabolome PCA of Cellvibrionaceae representatives.** *N. reynei* binned gills-symbiotic genomes gills.bin.1 and gills.bin.4 secondary metabolome grouped closely to a major group formed by *Teredinibacter* and influenced by BGCs for polyketide, non-ribosomal peptide, and hybrid compounds BGCs, besides putative routes for bacteriocins and terpenearylpolyene.

**Table S1 – Metagenomes overall features.** * Numbers (1,2,3): N. reynei specimens. Letters (A, B, C): metagenomic samples. ^1^ – QC – MG-RAST 4.0 applied quality control of the metagenomic paired end reads. 2 – Values obtained after metadata quality control pipeline filtering. Abbreviations: Int – Intestine; DG: Digestive Glands.

**Table S2 – CAZy domains detected on the binned genomes.**

**Table S3 – BLASTp analysis of** ***T. turnerae*** **T7901 multi-catalytic CAZymes using** ***N. reynei*** **symbiotic binned genome as database.** * T7901 CAZymes for which no highly conserved homolog could be detected on one or both genome bins. Stringency cutoff for highly conserved homologs are E value = 0.0, identity > 75% over the entire query protein.

**Table S4 – Genome bins contigs mapped to** ***Teredinibacter turnerae*** **T7901 putative Biosynthetic Gene Cluster.** * Characterized biosynthetic gene clusters for tartrolon antibiotics and turnerbactin siderophore production. Contigs highlighted in bold were detected by antismash server.

**Table S5: BLASTp analysis showing gills.bin.1 and gills.bin.4 BGCs conserved on** ***T. turnerae*** **species.**

**Table S6: BLASTp analysis of the genome bin gills.bin.4 unique BGCs core biosynthetic genes.**

**Table S7: Cellvibrionaceae representative genomes information and resistome and BGCs annotations.** * Coding sequence automated annotated by RAST server. † Resistance genes annotated according the CARD server and considering the default stringency cutoffs for perfect/strict and loosely annotations. ‡ Biosynthetic Gene Clusters (BGCs) annotated according to the antiSMASH server, under the bacterial version and using default settings.

